## Supplementary Information for "Intrinsically disordered SERBP1 regulates translation through topology-driven G-quadruplex recognition"

### Supplementary Tables

**Supplementary Table 1:** SERBP1 mutant constructs, mutations, and cloning primers used in the study.

| Construct | Mutations[a] | Addgene[b] | Forward primer | Reverse primer |
|---|---|---|---|---|
| SERBP1$^{\Delta RGG+\Delta C}$ | Δ369-400 | 253596 | CGTCCGGGTCGTGGTTAAGGATCCGGCTGCTAACAAAGCCCG | ACCACGACCCGGACGACCCAGATCACCGAAGTTGATTTCCAGCTGG |
| SERBP1$^{\Delta C}$ | Δ385-400 | 253597 | GGCCGTCCGAACCGTTAAGGATCCGGCTGCTAACAAAGCCCGAAAGG | ACGGTTCGGACGGCCACCACGACCACGGCCACCACGGCCACC |
| SERBP1$^{\Delta RGG}$ | Δ368-382 | 253598 | GGTCGTCCGGGTCGTAACCGTGGCAGCCGTACCGACAAG | ACGACCCGGACGACCCAGATCACCGAAGTTGATTTCCAGCTGGCTGG |
| SERBP1$^{189\text{-}400,S394G}$ | S394G | 253599 | AGCGCTGGCGCTCCGGATGTTGATGATTAAGGATCCGGCTGC | GAGCGCCAGCGCTGCTCTTGTCGGTACGGCTGCCAC |
| SERBP1$^{189\text{-}400,S386E}$ | S386E | 253600 | CGTGGCGAACGTACCGACAAGAGCAGCGCTAGCGCTCCGG | GGTACGTTCGCCACGGTTCGGACGGCCACCACGACCACG |
| SERBP1$^{189\text{-}400,2SE}$ | S386E, S394E | 253601 | AGCGCTGAAGCTCCGGATGTTGATGATTAAGGATCCGGCTGC | GAGCTTCAGCGCTGCTCTTGTCGGTACGTTCGCCAC |
| SERBP1$^{189\text{-}400,4SE}$ | S386E, S391E, S392E, S394E | 253602 | GAAGAAGCTGAAGCTCCGGATGTTGATGATTAAGGATCC | TTCAGCTTCTTCCTTGTCGGTACGTTCGCCACG |

[a] Mutations are in refence to the C-terminal construct (SERBP1189-400) described in doi: 10.1007/s12104-021-10046-3
[b] Addgene accession numbers (https://www.addgene.org).

**Supplementary Table 2:** Nucleotide sequences of the G-quadruplexes used in the study.

| Oligonucleotide | Sequence |
|---|---|
| 7mer | GCGCGGG |
| $mTOR_{23}$ | CGGGGAAGGCGGGCGGUGGGGCAGGGGCC |
| $HDAC3_2$ | GGCGGCGGAGG |
| rAGA | AAAAAAAAAAAAAAAGGGGGAAAAAAAAAAAAAAA |
| TPG4 | CUAGCAGCUAUGGGGGAGCUGGGGAAGGUGGGAAU |
| Pu20m2 | TAAGGGAGGGCGGGAGGGAA |
| rPu20m2 | UAAGGGAGGGCGGGAGGGAA |

**Supplementary Table 3:** G-quadruplex sequences identified in the mTOR mRNA 5' UTR by RG4-seq.

| RG4 id | Chromosome | RG4 start | RG4 end | Transcripts | Sequence |
|---|---|---|---|---|---|
| 31153 | 1 | 11262507 | 11262537 | ENST00000361445 | GACAGCGGGGAAGGCGGGCGGTGGGGCAGG |
| 31154 | 1 | 11262509 | 11262539 | ENST00000361445 | AGGACAGCGGGGAAGGCGGGCGGTGGGGCA |

| RG4 id | Length | Technique | Cell Line | Species | Reference PMID[a] |
|---|---|---|---|---|---|
| 31153 | 30 | RG4-seq | HeLa | *Homo Sapiens* | 27571552 |
| 31154 | 30 | RG4-seq | HeLa | *Homo Sapiens* | 27571552 |

[a] PubMed Identifier (https://pubmed.ncbi.nlm.nih.gov)

### Supplementary Figures

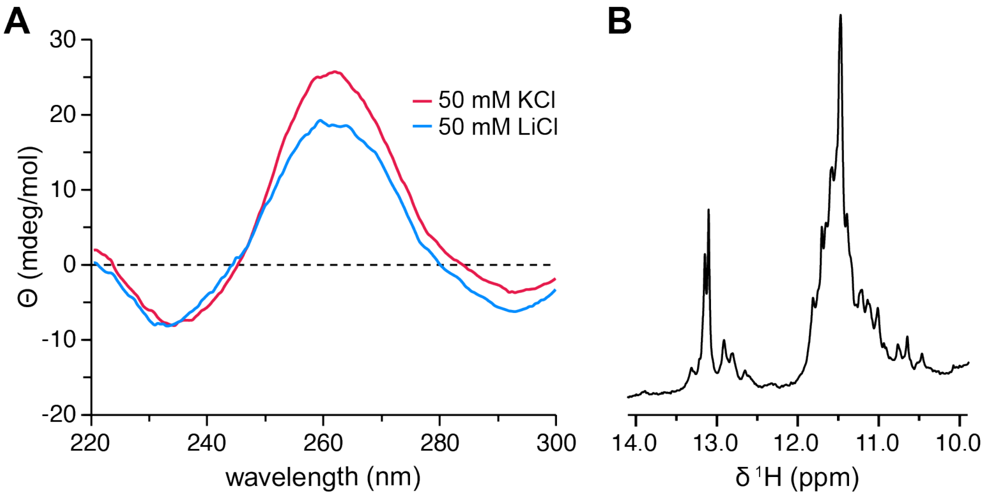


**Supplementary Figure 1: The 7mer oligonucleotide folds as a G-quadruplex.** (**A**) Circular dichroism spectrum of the 7mer in presence of $K^+$ (red) or $Li^+$ (blue). (**B**) Imino region of the $^1H$ NMR spectrum of the 7mer. Spectrum acquired at 278K on a 700 MHz spectrometer.

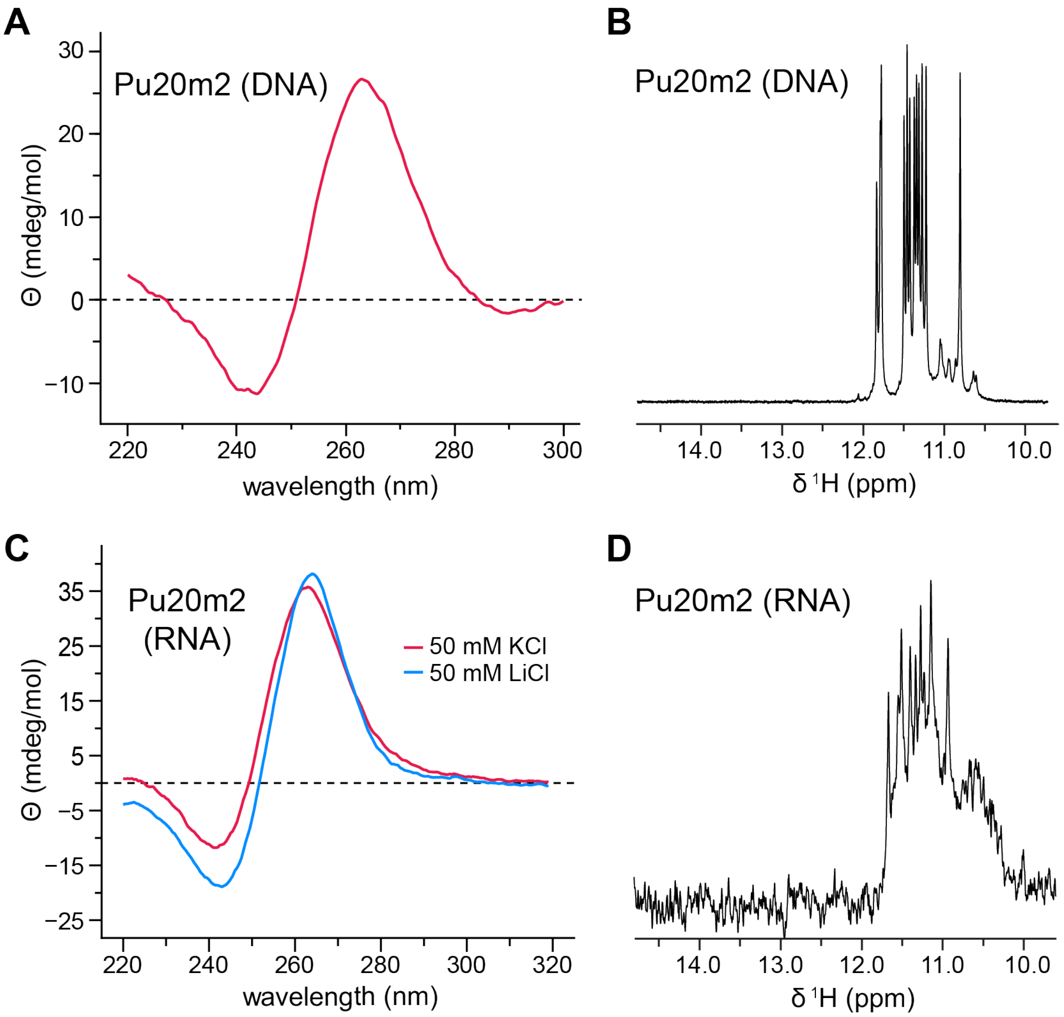


**Supplementary Figure 2: G4 folding of DNA and RNA Pu20m2.** (**A**,**C**) Circular dichroism spectrum of Pu20m2 and rPu20m2, respectively, in presence of $K^+$ (red) or $Li^+$ (blue). (**B**,**D**) Imino region of the $^1H$ NMR spectra of Pu20m2 and rPu20m2, respectively. Spectra were acquired on 1 mM (Pu20m2) and 100 µM (rPu20m2) samples at 278K on a 700 MHz spectrometer.

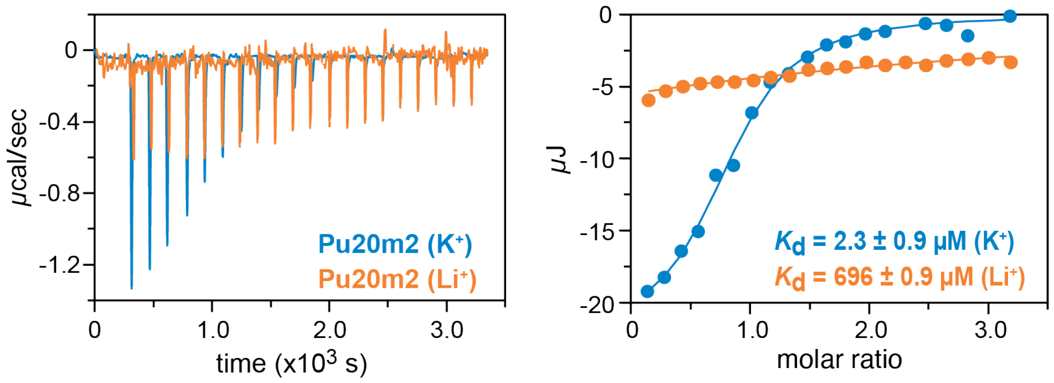


**Supplementary Figure 3: Isothermal titration calorimetry measurements of SERBP1$^{189\text{-}400}$ binding to Pu20m2.** Titrations were performed in the presence of either $K^+$ (blue) or $Li^+$ (orange).

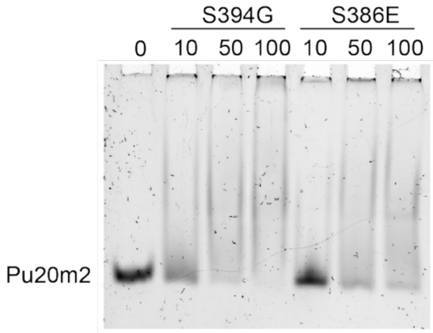


**Supplementary Figure 4: Electrophoretic mobility shift assay of Pu20m2 with SERBP1$^{189\text{-}400}$ point mutants.** SERBP1$^{189\text{-}400,\ S394G}$ and SERBP1$^{189\text{-}400,\ S386E}$ were tested at protein:G4 ratios of 0, 10:1, 50:1 and 100:1.

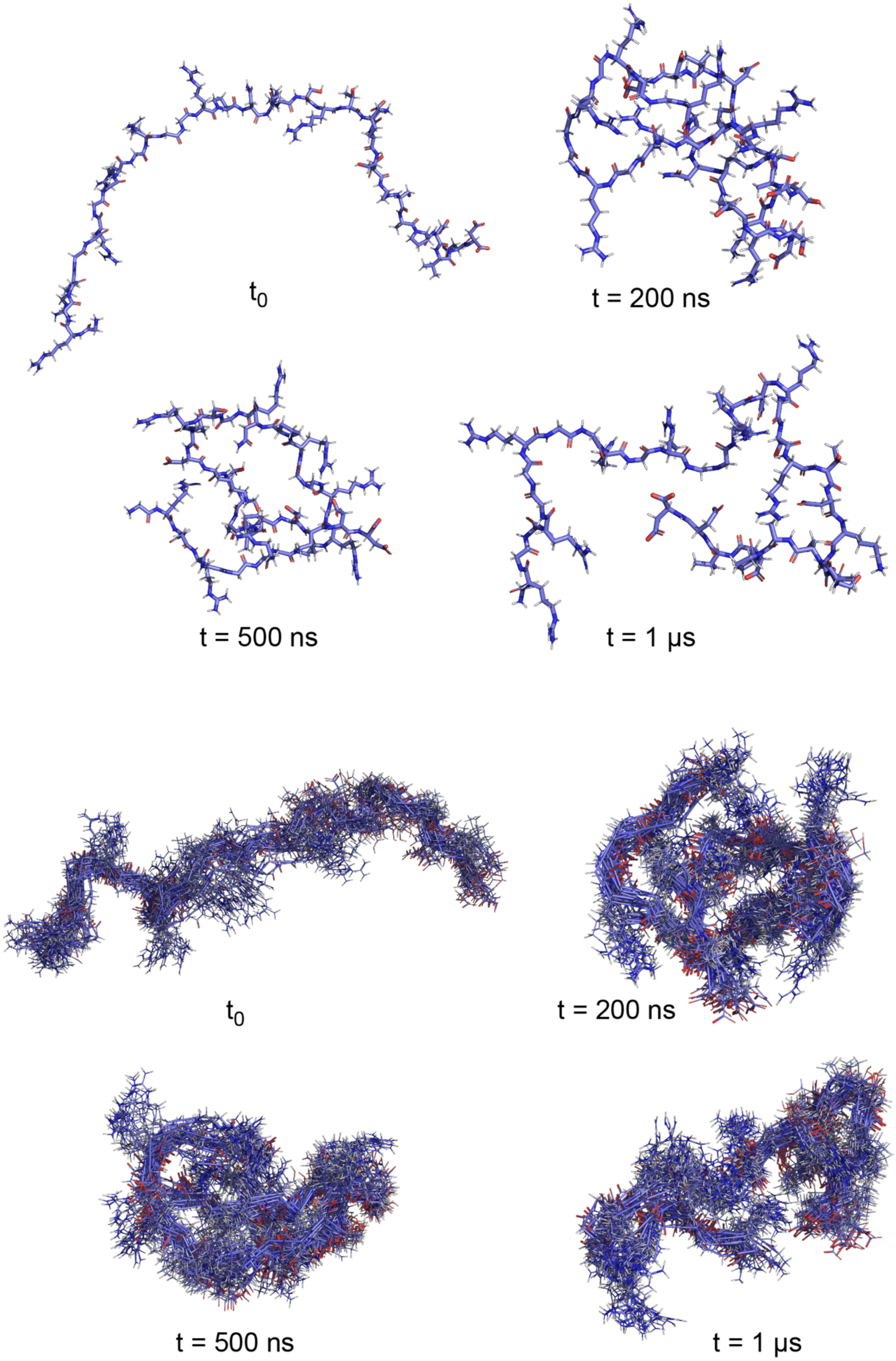


**Supplementary Figure 5: Snapshots of MD simulation for SERBP1$^{360\text{-}400}$**. Each ensemble is taken at different time points of the trajectory (0 ns, 200 ns, 500 ns, 900 ns) and is comprised of 20 consecutive structures spaced by 500 ps.

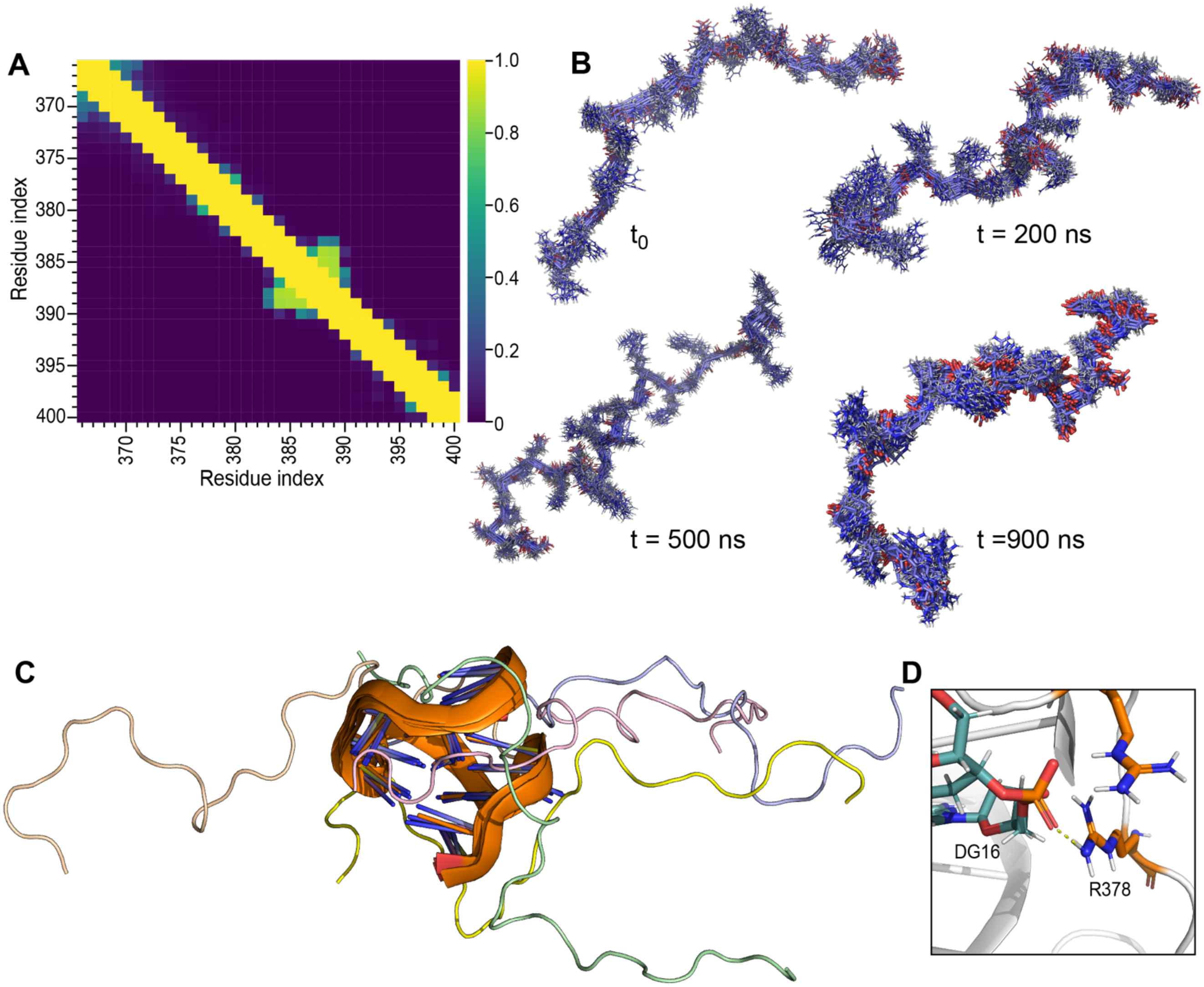


**Supplementary Figure 6: MD trajectory of SERBP1$^{360\text{-}400,4SE}$ and docking with Pu20m2.** (**A**) Contact map of SERBP1$^{366\text{-}400,4SE}$ residues with an 8.0 Å cutoff during MD simulation. Colors represent contact probability throughout the trajectory, from 0 (dark blue) to 1 (yellow). (**B**) Snapshots of the simulation at different time points (0, 200, 500, and 900 ns). (**C**) Superimposition of the five top-scoring SERBP1$^{366\text{-}400,4SE}$–Pu20m2 complexes from HADDOCK docking. (**D**) Atomic details of SERBP1$^{366\text{-}400,4SE}$–Pu20m2 interactions at the residue level.

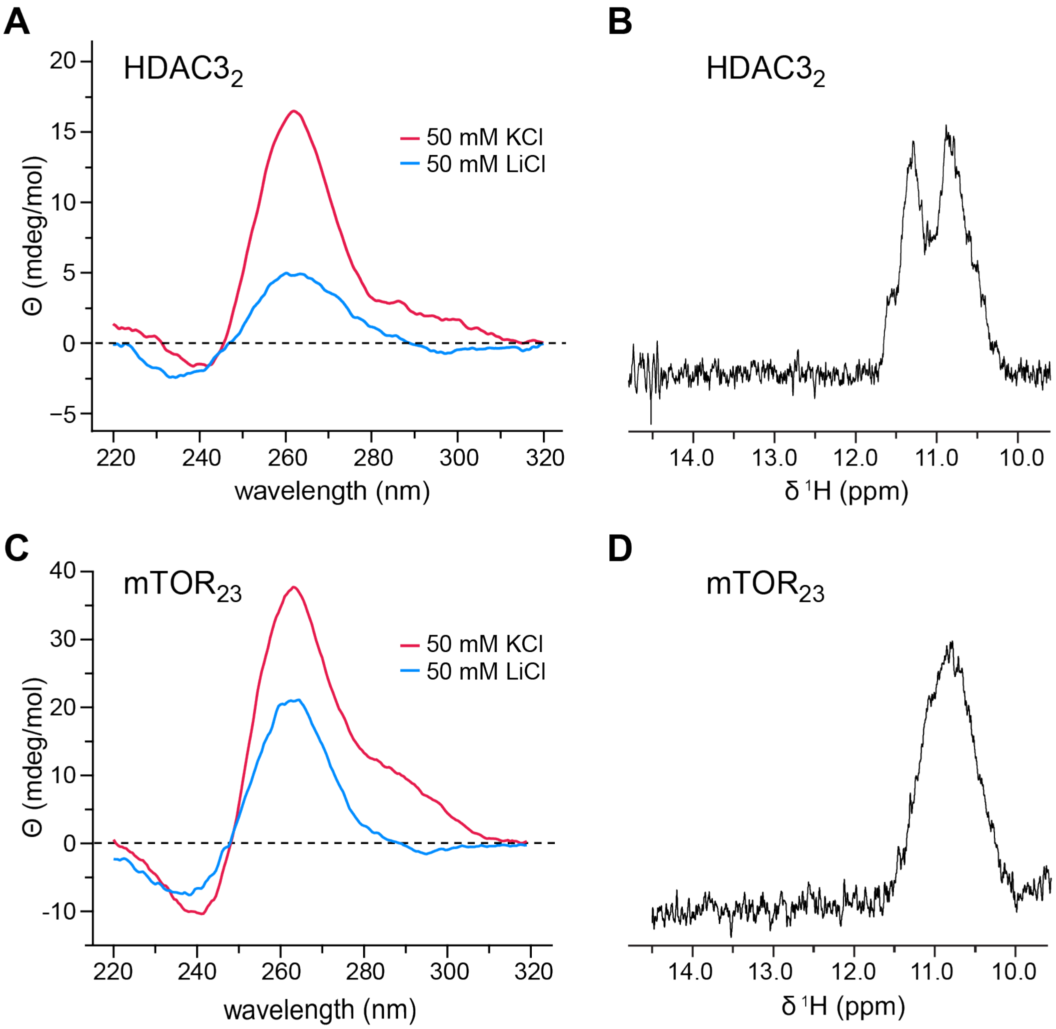


**Supplementary Figure 7: G4 folding of HDCA3$_2$ and mTOR$_{23}$ RNA sequences.** (**A**,**C**) Circular dichroism spectra of HDCA3$_2$ and mTOR$_{23}$, respectively, in presence of $K^+$ (red) or $Li^+$ (blue). (**B**,**D**) Imino region of the $^1$H NMR spectra of HDCA3$_2$ and mTOR$_{23}$, respectively. Spectra were acquired at 278K on a 700 MHz spectrometer.

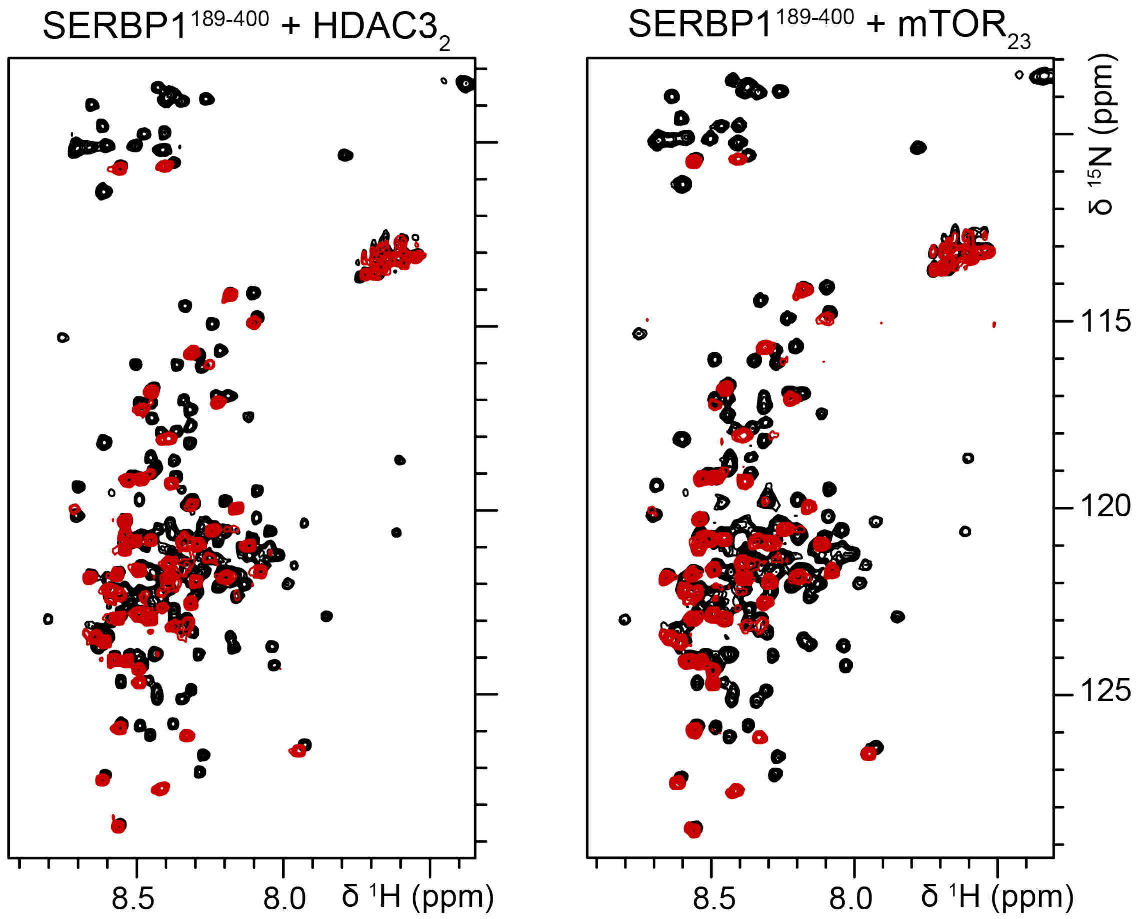


**Supplementary Figure 8: RNA G4 binding causes dramatic line broadening at the first titration point.** Overlay of $^{1}$H,$^{15}$N-HSQC spectra of SERBP1$^{189-400}$ recorded on protein alone (black) and at the first titration point (1:4 G4:protein ratio, red). Spectra were acquired at 278K on a 700 MHz spectrometer.
